# Regulation of DNA methylation on key parasitism genes of Cysticercus cellulosae revealed by integrative epigenomic-transcriptomic analyses

**DOI:** 10.1101/353417

**Authors:** Shumin Sun, Xiaolei Liu, Guanyu Ji, Xuelin Wang, Junwen Wang, Xue Bai, Jing Xu, Jianda Pang, Yining Song, Xinrui Wang, Fei Gao, Mingyuan Liu

## Abstract

**Background:** The life cycle of *Taenia solium* is characterized by different stages of development, requiring various kinds of hosts that can appropriately harbor the eggs (proglottids), the oncospheres, the larvae and the adults. Similar to other metazoan pathogens, *T. solium* undergoes transcriptional and developmental regulation via epigenetics during its complex lifecycle and host interactions.

**Result:** In the present study, we integrated whole-genome bisulfite sequencing and RNA-seq technologies to characterize the genome-wide DNA methylation and its effect on transcription of Cysticercus cellulosae of *T. solium*. We confirm that the *T. solium* genome in the cysticercus stage is epigenetically modified by DNA methylation in a pattern similar to that of other invertebrate genomes, i.e., sparsely or moderately methylated. We also observed an enrichment of non-CpG methylation in defined genetic elements of the *T. solium* genome. Furthermore, an integrative analysis of both the transcriptome and the DNA methylome indicated a strong correlation between these two datasets, suggesting that gene expression might be tightly regulated by DNA methylation. Importantly, our data suggested that DNA methylation might play an important role in repressing key parasitism-related genes, including genes encoding excretion-secretion proteins, thereby raising the possibility of targeting DNA methylation processes as a useful strategy in therapeutics of cysticercosis.

**Conclusion:** Our study will provide a foundation for future studies to explore this key epigenetic modification in development of Cysticercus cellulosae and in human cysticercus disease.

## Introduction

Cysticercus cellulosae, the larval stage of *T. solium*, resides in the central nervous system, skeletal muscle, and other organs of both pigs and humans [1], resulting in the high prevalence of cysticercosis worldwide. As a neglected tropical disease prioritized by the World Health Organization, serious human disease burden [2] and annual economic losses in livestock are caused by infection with this pork tapeworm. To better control this disease, the mechanisms of transcriptional and developmental regulation during its complex lifecycle and host interactions should be better understood.

ADNA methylation, i.e., 5-methylcytosine (m5C) is an important epigenetic mechanism that is present in the genomes of *Trichinella spiralis* [3] and Platyhelminthes (*Schistosoma mansoni* [4]) parasitic nematodes. Via regulating gene transcription, DNA methylation plays an important role in parasitism. Similar to *T. solium*, *S. mansoni* belongs the phylum of Platyhelminthes. A previous study showed that *S. mansoni* contains conserved DNA methyltransferase 2 (DNMT2) and methyl-CpG binding proteins (MBD) [4]. Importantly, demethylation induced by 5-azacytidine can disrupt egg production and maturation, indicating an essential role for DNA methylation in the normal development of this parasitic worms in this phylum [4].

In the present study, we aimed to identify functional DNA methylation machinery and detect cytosine methylation levels in the cysticercus cellulosae of *T. solium* based on a draft genome that has been sequenced and annotated previously [5]. To achieve this aim, we applied the whole-genome bisulfite sequencing (WGBS) method to characterize the genome-wide DNA methylation pattern at single-base resolution [6]. Based on this unbiased characterization, our results confirm that in the cysticercus stage, the *T. solium* genome [5] is epigenetically modified by DNA methylation in a pattern similar to that of other invertebrate genomes, i.e., sparsely or moderately methylated [7, 8]. We also observed an enrichment of non-CpG methylation in defined genetic elements of *T. solium* genome, which is a pattern different from mammalian methylomes [7, 8]. Furthermore, we applied RNA-seq technology to profile gene expression. An integrative analysis on both the transcriptome and DNA methylome indicated a strong correlation between these two datasets, suggesting that gene expression might be tightly regulated by DNA methylation. Importantly, our data suggested that DNA methylation might play an important role in repressing key parasitism-related genes, including genes encoding excretion-secretion proteins. In summary, for the first time, we provide data to characterize the DNA methylome and the transcriptome of the *T. solium* cysticercus cellulosae. Our data will be valuable to the community and will allow researchers to provide new insights into the mechanism of methylation in cysticercosis in future studies.

## Materials and Methods

### Sample collection and nuclei acid extraction

Individual cysticerci were isolated from a single, naturally infected pig (Neimeigu Province, China) and rinsed thoroughly several times with phosphate-buffered saline. The cysticerci were first frozen in liquid nitrogen and then finely ground to a powder-like texture. Genomic DNA was extracted using the phenol chloroform extraction method, and total RNA was purified using Trizol reagent (Invitrogen, CA, USA) according to the manufacturer’s instructions. RNA was dissolved in diethylpyrocarbonate (DEPC)-treated water and treated with DNase I (Invitrogen, CA, USA). The quantity and quality of the DNA and RNA were tested by ultraviolet-Vis spectrophotometry with a NanoDrop 2000 (Thermo Scientific CA, USA).

### BlastP searches and phylogenetic analysis of DNMTs

Reciprocal BlastP comparisons were first performed to identify DNMTs and MBD orthologs. Significant hits were defined as those satisfying the following criteria: E-value < 1e-5 and aligned segments covering at least 30% of the sequence length of the hit. For phylogenetic analysis, multiple sequence alignment was performed by Clustal W [9]. The MEGA7 with the neighbor-joining method [10, 11] based on the JTT+ G (Jones-Taylor-Thornton and Gamma Distribution) model was applied to reconstruct the phylogenetic tree.

### MethylC-seq library construction and sequencing

Prior to library construction, 5 μg of genomic DNA extracted from a cysticercosis body was spiked with 25 ng unmethylated lambda DNA (Promega, Madison, WI, USA) and fragmented using a Covarias sonication system to a mean size of approximately 200 bp. After fragmentation, libraries were constructed according to the Illumina Paired-End protocol with some modifications. Briefly, purified randomly fragmented DNA was treated with a mix of T4 DNA polymerase, Klenow fragment and T4 polynucleotide kinase to repair blunt ends and phosphorylate the ends. The blunt DNA fragments were subsequently 3’ adenylated using Klenow fragment (3’-5’ exo-), followed by ligation to adaptors synthesized with 5’-methylcytosine instead of cytosine using T4 DNA ligase. After each step, DNA was purified using a QIAquick PCR purification kit (Qiagen, Shanghai, China). Next, a ZYMO EZ DNA Methylation-Gold Kit^TM (ZYMO Research, Irvine, CA, USA) was employed to convert unmethylated cytosine to uracil, according to the manufacturer’s instructions, and 220 to 250 bp converted products were size selected. Finally, PCR was carried out in a final reaction volume of 50 μl consisting of 20 μl of size selected fractions, 4 μl of 2.5 mM dNTPs, 5 μl of 10× buffer, 0.5 μl of JumpStart™ Taq DNA Polymerase, 2 μl of PCR primers and 18.5 μl water. The thermal cycling program was 94°C for 1 minute; 10 cycles of 94°C for 10 s, 62°C for 30 s, 72°C for 30 s; and then a 5-minute incubation at 72°C before holding the products at 12°C. The PCR products were purified using a QIAquick gel extraction kit (Qiagen). Before analysis with an Illumina *Hiseq2500*, the purified products were analyzed using a Bioanalyzer analysis system (Agilent, Santa Clara, CA, USA) and quantified by real-time PCR. Raw sequencing data were processed using the Illumina base-calling pipeline (Illumina Pipeline version 1.3.1). The sodium bisulfite non-conversion rate was calculated as the percentage of cytosines sequenced at cytosine reference positions in the lambda genome.

### RNA-seq library construction and sequencing

Total RNA was extracted using the Invitrogen TRIzol Reagent and then treated with RNase-free DNase I (Ambion, Guangzhou, China) for 30 minutes. The integrity of total RNA was checked using an Agilent 2100 Bioanalyzer. cDNA libraries were prepared according to the manufacturer’s instructions (Illumina). The poly(A)-containing mRNA molecules were purified using Oligo (dT) Beads (Illumina) and 20 μg of total RNA from each sample. Tris-HCl (10 mM) was used to elute the mRNA from the magnetic beads. To avoid priming bias when synthesizing the cDNA, mRNA was fragmented before cDNA synthesis. Fragmentation was performed using divalent cations at an elevated temperature. The cleaved mRNA fragments were converted into double-stranded cDNA using SuperScript II, RNase H and DNA Pol I, primed by random primers. The resulting cDNA was purified using a QIAquick PCR Purification Kit (Qiagen). Then, the cDNA was subjected to end repair and phosphorylation using T4 DNA polymerase, Klenow DNA polymerase and T4 Polynucleotide Kinase (PNK). Subsequent purifications were performed using the QIAquick PCR Purification Kit (Qiagen). These repaired cDNA fragments were 3’-adenylated using KlenowExo (Illumina) and purified using the MinElute PCR Purification Kit (Qiagen), producing cDNA fragments with a single ‘A’ base overhang at the 3’ end for subsequent ligation to the adapters. Illumina PE adapters were ligated to the ends of these 3’-adenylated cDNA fragments and then purified using the MinElute PCR Purification Kit (Qiagen). To select a size range of templates for downstream enrichment, the products of the ligation reaction were purified on 2% TAE-Certified Low-Range Ultra Agarose (Bio-Rad, Hercules, CA, USA). cDNA fragments (200 ± 20 bp) were excised from the gel and extracted using the QIAquick Gel Extraction Kit (Qiagen). Fifteen rounds of PCR amplification were performed to enrich the adapter-modified cDNA library using primers complementary to the ends of the adapters (PCR Primer PE 1.0 and PCR Primer PE 2.0; Illumina).

### Transcriptome mapping

RNA-seq reads were trimmed to a maximum length of 80 bp, and stretches of bases having a quality score <30 at the ends of the reads were removed. Reads were mapped using Tophat 2.0.11 [12]. As reference sequence for the transcriptome mapping we used the current assembly of the *T. solium* database [13]. Expression was quantified using cufflinks 2.1.1 [14]. RepeatMasker [15] were used to identify tandem repeats.

### Bisulfite mapping and methylation calling

Reads were trimmed to a maximal length of 125 bp, and stretches of bases having a quality score <30 at the ends of the reads were removed. Reads were mapped using BSMAP 2.2.74 [16]. As a reference sequence for the bisulfite mapping we used the current assembly of the *T. solium* genome [13]. Only reads mapping with both partners of the read pairs at the correct distance were used. The CpG-specificity was calculated by determining the number of cytosines called in all mapped reads at all non-CpG positions and dividing by the number of all bases in all mapped reads at all non-CpG positions. Methylation ratios were determined using a Python script (methratio.py) distributed together with the BSMAP package for both the forward and reverse strands.

### Protein network analyses

The STRING online tool [17] was used with default parameters. Peptide sequences of key genes were the input and were aligned to *Caenorhabditis elegans* protein sequences.

### Data availability

The *T. solium* methylome data have been deposited at NCBI/GEO/ under the accession number GSE84086.

## Results

### The presence of DNA methylation in the *T. solium* genome

The methylation status of DNA is related to three types of enzymes, including DNA methyltransferases, which affect maintenance methylation and de novo methylation. To understand whether *T. solium* possesses the ability to methylate DNA, we first conducted a reciprocal Blast alignment to identify genes that might be homologous to known DNA (cytosine-5)-methyltransferases. As a result, two genes (Scaffold00200.gene8095 and LongOrf.asmbl_16366) were identified that are homologous to *DNMT3B* and *DNMT2*, respectively, with high sequence similarity (e-value < 1e-10). Scaffold00067.gene4890 was aligned (e-value < 1e-5) with either *DNMT3A* or *DNMT3B* from multiple species. In addition, more than one gene was matched with *DNMT1*, among which Scaffold00068.gene4920 had the best hit (e-value < 1e-10) (Table S1). Phylogenetic analyses by MEGA7 also supported these results (Figure S1). Moreover, we searched for genes homologous to methyl-CpG binding domain protein (MBD). Two candidate genes (LongOrf.asmbl_5021 and LongOrf.asmbl_14047) were homologous to *MBDs* in multiple species, including *Echinococcus granulosus*, which is closely related to *T. solium* evolutionarily (Table S1 and Figure S2). A full repertoire of functionally conserved amino acid residues was identified for both the potential DNMT2 and DNMT3 and the MBDs of *T. solium*, indicating that these proteins are functionally active (Table S2). However, a high level of divergence between *T. solium* and other species was observed for DNMT1 homologs (Table S2), which was in agreement with previous studies.

Given these results, we assessed the genome-wide DNA methylation profiles in *T. solium* using MethylC-Seq. There were 54.21 million raw reads generated (Table S3). BSMAP [16] was used to align the sequenced reads to the *T. solium* reference sequence, reaching an approximately 76.41% mapping rate. The average read depth was 11.32 per strand, while on average, over 50 Mb (90.52%) of each strand of the *T. solium* reference sequence was covered. Because of the potential occurrence of non-conversion and thymidine-cytosine sequencing errors, the false-positive rate was estimated by calculating the methylation level of lambda DNA, which is normally unmethylated (Materials and methods). We then applied the error rate (0.0041) to correct methylated cytosine sites (mC) identification according to the method described by Lister et al. [6], which is based on a binomial test and false discovery rate constraints. As a result, approximately 76.6 thousand mCs were estimated in the *T. solium* genome (accounting for 0.20% of the total cytosines sequenced with depth ≥5X). Both symmetrical CpG methylation and asymmetrical non-CpG methylation were revealed.

### Characterization of overall methylation patterns

We further characterized the global patterns of DNA methylation in the genomes of *T. solium*. First, we showed the percentage of methylated cytosine of each sequence context. Among the 76.6 thousand mCs across the entire genome, a majority (69.5%) were in the context of CHH. In contrast, only 15.38% and 15.12% of the mCs were located in the contexts of CHG and CpG, respectively (Figure S3 A). Furthermore, most of the CpG and non-CpGs displayed a low methylation fraction (<30%) (Figure S3B and C). These patterns are highly different from mammalian methylomes, in which most 5mCs are located in CpG contexts and the majority of the CpGs are highly methylated (>50%) [7]. Since most (69.5%) of the mCs in the *T. solium* genome were in the CHH context, we further analyzed the sequence context of mCHHs across the entire genome to further examine whether there is any sequence bias in the enrichment of cytosine methylation in the CHH context. As a result, mCpA was shown to be preferentially enriched within the methylated CHH dinucleotide (Figure 1A). There were more than 21,000 methylated CpAs in each strand, meaning that 55.73% of total CpAs were methylated in the entire genome (Figure 1B, D). This result was consistent with reports that mCpA was predominantly found in another tapeworm, *S. mansoni* [18, 19]. With regard to methylation levels, we did not observe significant differences among different sequence contexts for mCs (Figure 1C).

**Figure 1.**
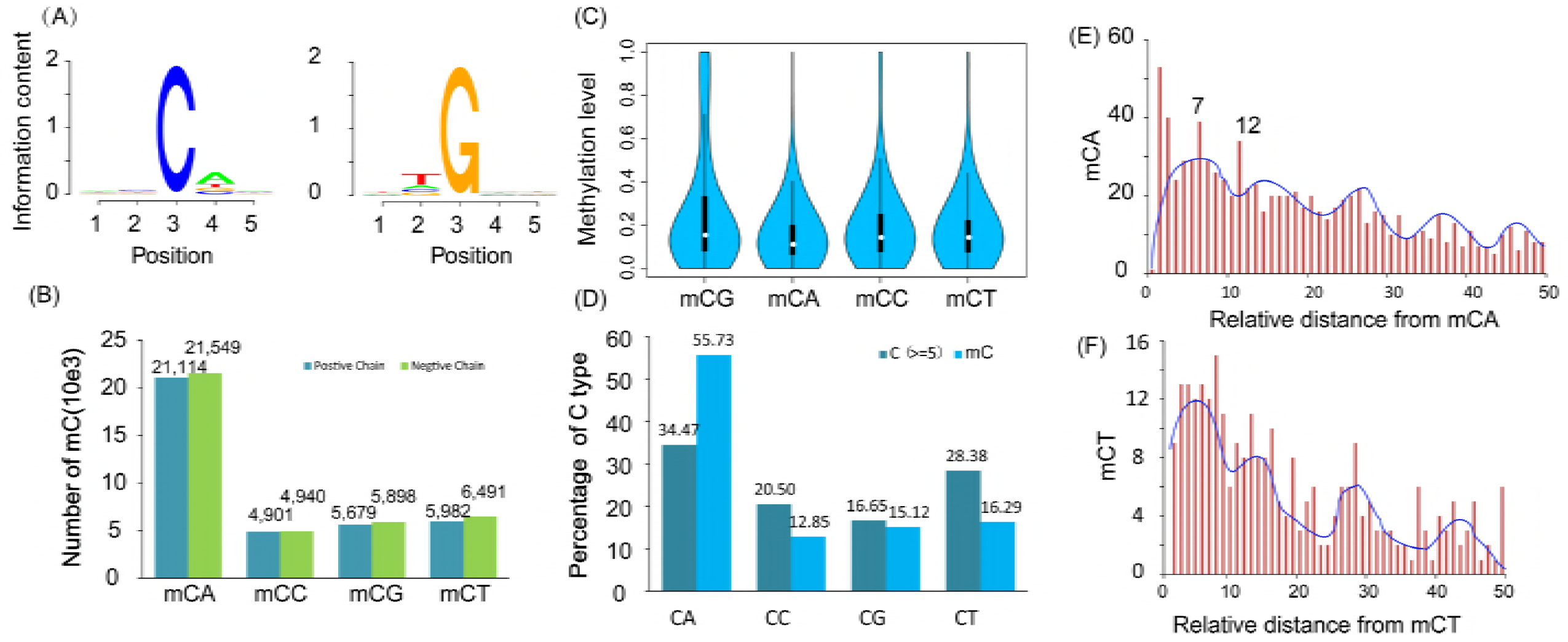
Cytosine DNA methylation in *T. solium*. (A) Logo plots of the sequences proximal to sites of cytosine DNA methylation in each sequence context in *T. solium*; (B) Number of mCs for each type of dinucleotide; (C) Distribution of mCs; (D) Percentage of each type of dinucleotide; (E, F) Prevalence of mCA/mCT sites (y-axis) as a function of the number of bases between adjacent mCA/mCT sites (x-axis) based on all non-redundant pair-wise distances up to 50 nt in all introns. The blue line represents smoothing with cubic splines.

We next examined whether there was any preference for the distance between adjacent sites of DNA methylation in the *T. solium* genome. The relative distance between mCs in each context within 50 nucleotides in introns was then analyzed because of the steady methylation without any selective pressure by protein coding genes in intron regions. Similar to the periodicity of 8-10 bases revealed in previous studies on the Arabidopsis and human genomes [20], we also observed a strong tendency of peaked enrichment of mCpA sites, which might be explained by a single turn of the DNA helix (Figure 1E). Moreover, we found that mCpT revealed a similar periodicity of 8-12 bases (Figure 1F), though the numbers of cytosines in the context of CpG and CpC were too few to yield reliable results (Figure S3D and E). In summary, our results indicated that the molecular mechanisms governing de novo methylation at CpA sites may be similar among the *cysticercus* and the plant and animal kingdoms.

We then examined the distribution of methylation levels for the four categories of methylated cytosines across the entire genome. In general, similar mosaic distribution patterns were observed for methylation levels of all types of mCs, that is, relatively highly methylated domains were interspersed within regions with low methylation (Figure S4A). Furthermore, the distribution of mCs across the genome was also uneven; dense mCs of specific categories were occasionally enriched in specific scaffolds (Figure S4B). Such a pattern has been observed in previous studies on other invertebrates. We also examined the patterns of methylation in annotated elements, including genes, tandem repeats, and transposable elements. The methylation percentage of each cytosine context in exons was higher than that in other annotated elements, especially CpAs, which accounted for a more than 2-fold greater percentage than the other contexts in exons (Figure 2A).

**Figure 2.**
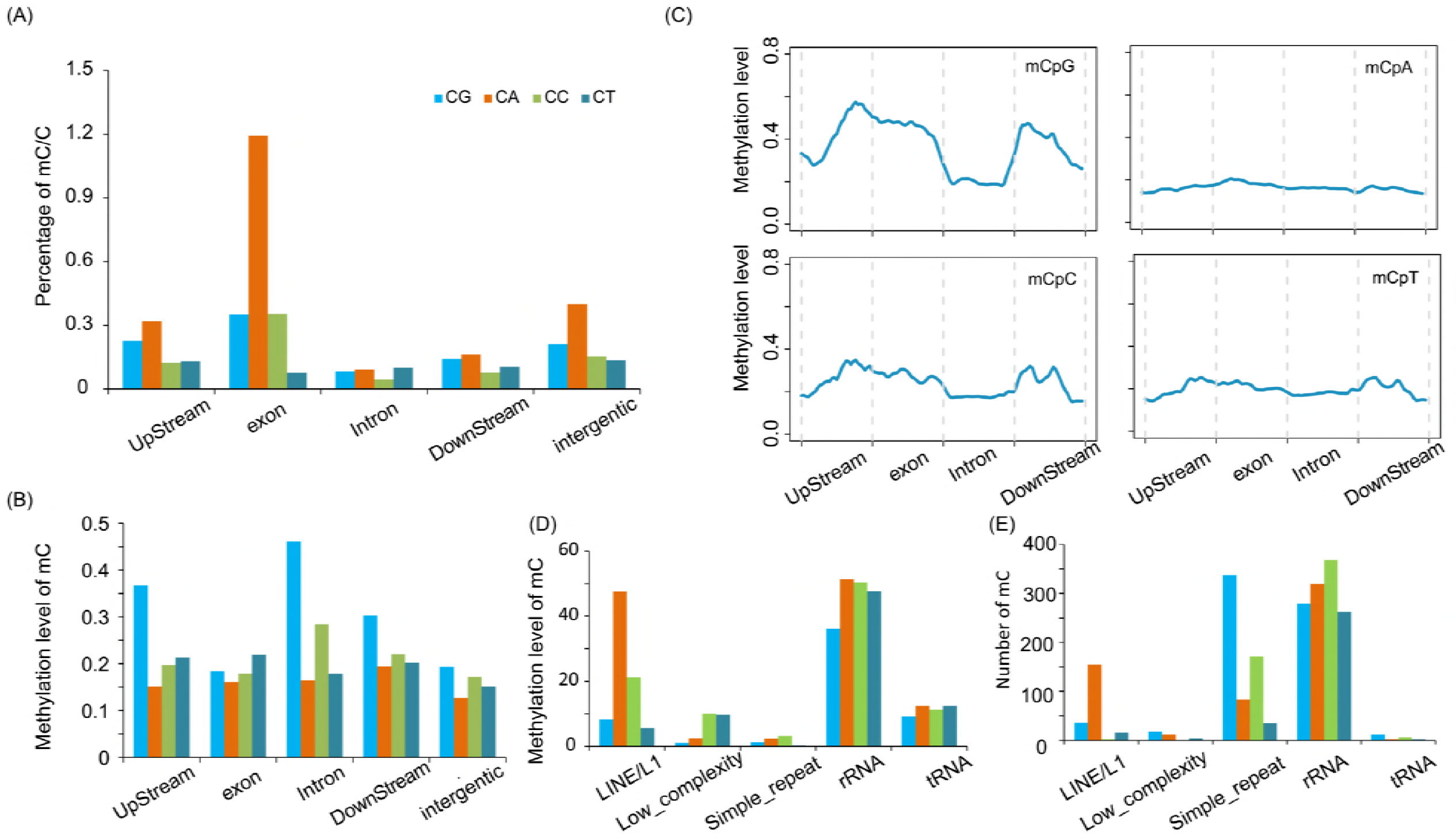
Average methylation levels of different genomic regions. (A, B) Average density of methylation; (C) Average density of mC methylation distributed on the genome. Two-kilobase regions upstream and downstream of each gene were divided into 100-bp (bp) intervals. Each coding sequence or intron was divided into 20 intervals (5% per interval). (D) Average mC methylation level on repeat elements; (E) Number of mCs on each repeat element.

We then examined the average methylation level in each element, which showed that average CpG methylation levels were higher than those other types of methylated cytosines, similar to mammalian genomes. However, the genome-wide pattern was again divergent from mammalian genomes, as higher average methylation in exons and lower methylation in introns of CpG sites were observed (Figure 2B, C and Figure S5). The trend for the average methylation of CpC and CpT was similar to that of CpG. However, a uniform distribution of CpA methylation levels in each annotated element was displayed (Figure 2C and Figure S5). We also analyzed the methylation level of each cytosine context in repeat regions (Figure 2D, E). Previous studies have indicated that transposable elements are usually unmethylated in the honey bee Apis mellifera and silkworm Bombyx mori [21, 22]. In cysticercus, we observed a similar phenomenon as the above species except that relatively highly methylated rRNAs were observed in *T. solium*. Notably, CpAs were methylated at a higher level or frequency than other types in LINE/L1 (Figure 2D, E).

### The relationship between methylation and gene expression

It was reported that DNA methylation plays an important role in regulating gene expression. We evaluated gene expression in *T. solium* using Illumina high-throughput RNA-seq technology. Most of the raw reads could be uniquely mapped to previously annotated genes (88.17%). A total of 9,718 annotated genes out of 11,903 could be aligned with at least one unique read. To characterize the relationship between DNA methylation and gene expression, we divided the expressed genes with at least one read into quartiles of expression levels. We then examined the distribution of methylation levels for different quartiles of expressed genes and genes exhibiting no expression. High CpG and CpC methylation levels were observed in upstream and exon regions of genes with the lowest expression. Moreover, a negative correlation could also be observed between CpA and CpT methylation levels of upstream and exons and expression levels of these expressed genes. However, for silent genes, mainly high CpG and CpC methylation levels of downstream regions were observed (Figure 3). Taken together, methylation levels of mCs from both CpG or non-CpG sequence contexts were correlated with gene expression levels, though different regulation mechanisms might be involved.

**Figure 3.**
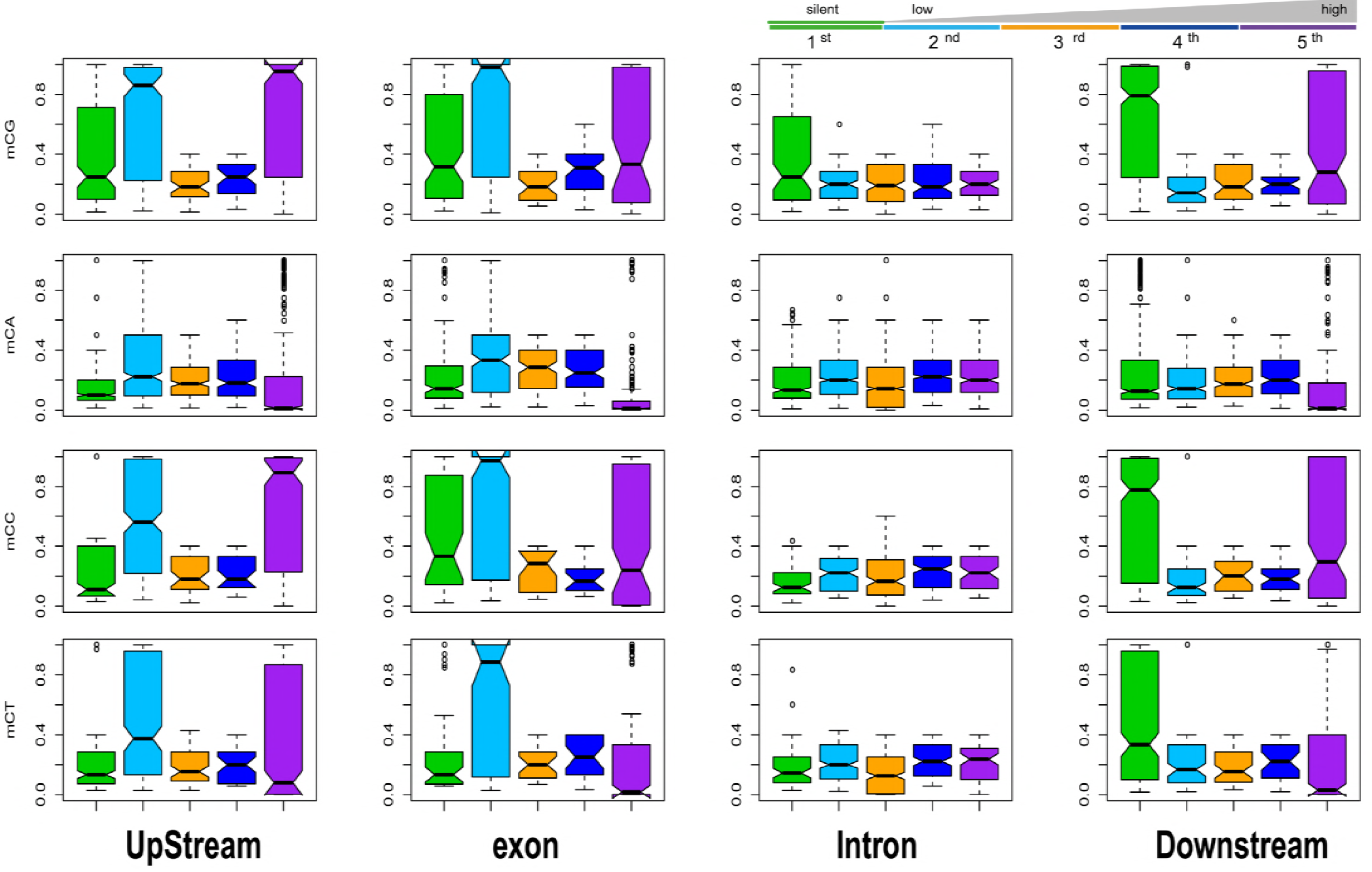
Relationship between mC DNA methylation and expression levels of genes in *T. solium*. Percentage of methylation within genes that were classified based on expression levels. The first class includes silent genes with no sequencing reads detected, and the second to fifth classes cover expressed genes from the lowest 25% to the highest 25%. Regions of 2 kb upstream and downstream of each gene was divided into 100-bp intervals, and each gene was divided into 20 intervals (5% per interval).

Next, to infer whether methylated genes were enriched for specific molecular functions, we filtered out a total of 1,647 of the genes with the lowest expression and 1,354 of the most highly expressed genes, based on the criteria that at least one mC was present within their genic regions. Then, we applied the WEGO (Web Gene Ontology Annotation Plotting) tool [23] to functionally categorize the gene ontology (GO) terms of these genes. We found that these two sets of genes displayed similar patterns of GO enrichment, specifically, “cell” and “cell part” in Cellular Component, “binding” and “catalytic” Molecular Functions, and “cellular process” and “metabolic process” in Biological Process were relatively enriched. This result suggested that the genes heavily regulated by DNA methylation were more prone to signaling regulation or interaction with environmental factors, e.g., diet or metabolism (Figure S6). In summary, these results suggested the potential for the regulation of *T. solium* genes by DNA methylation, especially those that function as regulators of cell-cell or cell-environmental communication. Furthermore, different molecular mechanisms might be involved depending on different mC contexts and genes.

### Regulation of DNA methylation on key parasitism genes of *T. solium*

To obtain further insight into the epigenetic regulation of parasite development, survival and parasite-host interactions of *T. solium*, we next studied conserved genes across tapeworm-species and genes encoding excretion–secretion proteins (ESPs) in *T. solium*. For conserved genes, we applied a gene set that was reported previously in a study by Bjorn Victor et al., in which 261 genes conserved between Taenia and Echinococcus tapeworms were obtained by comparing the transcriptomes of five important intestinal parasites, including *T. multiceps*, *T. solium*, *E. granulosus*, E. multilocularis and *T. pisiformis* [24]. Based on their results, we further retrieved 216 genes with the best blastx hit for each contig (e<1e-10) and studied their DNA methylation status. A total of 190 of these genes contained at least one mC across their genic regions. As indicated in Figure 4C, CpG and CpC methylation levels in upstream and exon regions were higher than other types of methylation and in other genic regions. A further examination of the 190 genes revealed that 71 genes contained CpG or CpC methylation within their upstream or exon regions. Therefore, we searched for extensively methylated genes based on the criterion that the CpG and CpC methylation levels of the examined gene were significantly higher than the average value of the 71 genes. As a result, we revealed 14 conserved genes that were extensively methylated on CpG/CpC sites within their upstream regions and exons (p<0.05). Compared with those 26 genes without mC, we found these 14 genes were expressed at a significantly lower level (Figure 4A), suggesting DNA methylation is a key mechanism for the transcriptional regulation of these conserved genes. For ESPs, we also applied a dataset containing 76 ESPs for *T. solium*, which was identified by Bjorn Victor et al. using a proteomics strategy [25]. We applied the BlastP algorithm to align these ESPs back to the genome and revealed 111 gene sequences that might encode these ESPs (Table S4). Using the same criterion for conserved genes, we found 13 extensively methylated genes. Similarly, gene expression comparisons again revealed that these 13 genes were expressed at significantly lower levels than the 26 non-methylated genes (Figure 4A). Using a similar strategy, we also looked into genes containing methylated CpAs and CpTs within their upstream or exon regions. However, no clear difference in gene expression levels was observed (Figure 4B). These results indicated that CpG/CpC methylation in upstream regions and exons played a major role in *T. solium* gene repression. Furthermore, we found a different distribution pattern between mCpG and mCpC for these repressed genes, in which mCpG were mostly distributed in upstream regions, while mCpC were more often in exons (Figure S5). Based on the above analyses, we revealed 27 key genes that might be repressed by DNA methylation mechanisms. Interestingly, a protein-protein interaction analysis using the STRING online tool [17] indicated strong mutual interactions among these conserved proteins and ESPs (Figure 5) based on annotation of the model organism *Caenorhabditis elegans*.

**Figure 4.**
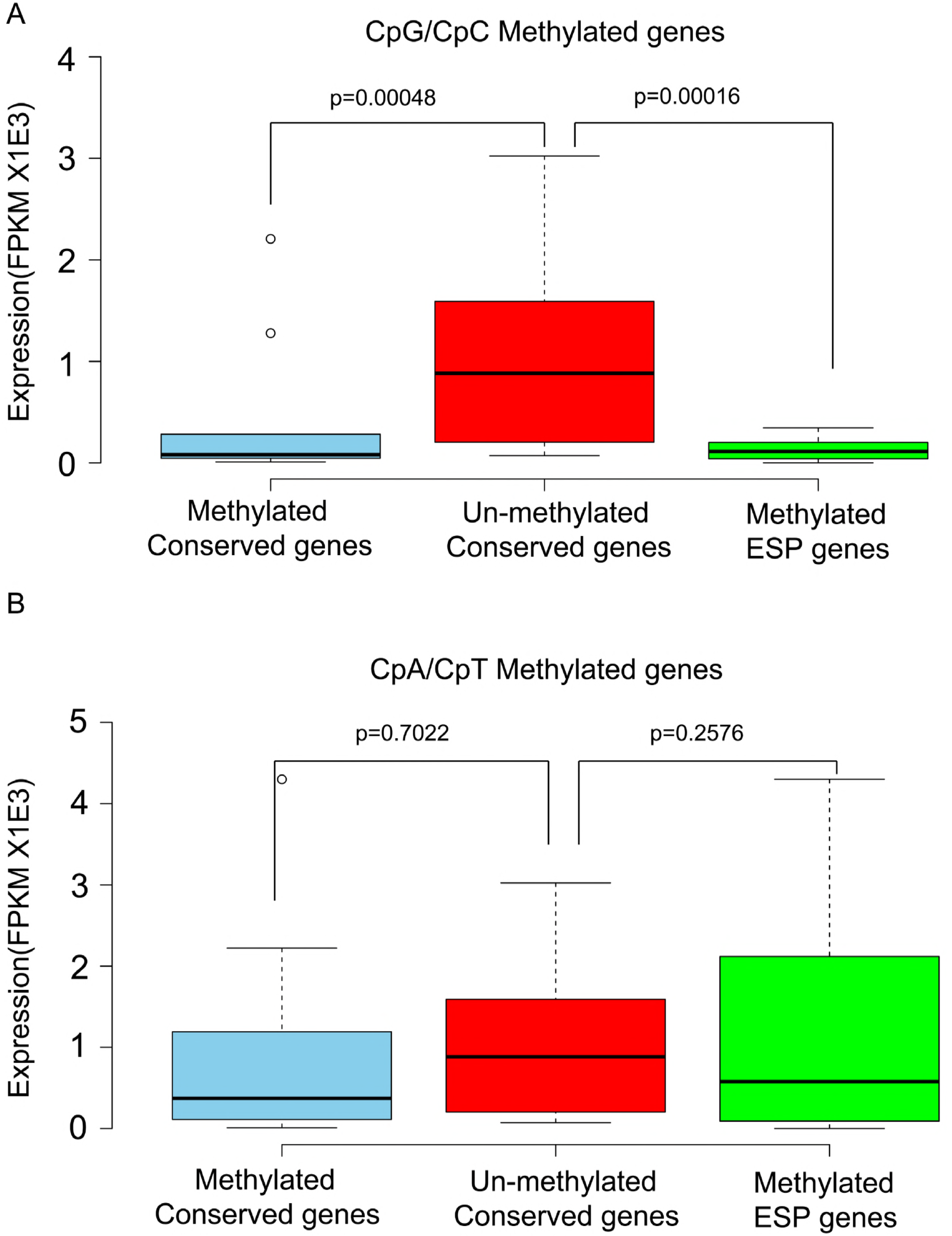
Boxplots of gene expression levels of un-methylated genes, and genes with conserved methylation and methylated ESP genes based on either (A) CpG/CpC methylation or (B) CpA/CpT methylation.

**Figure 5.**
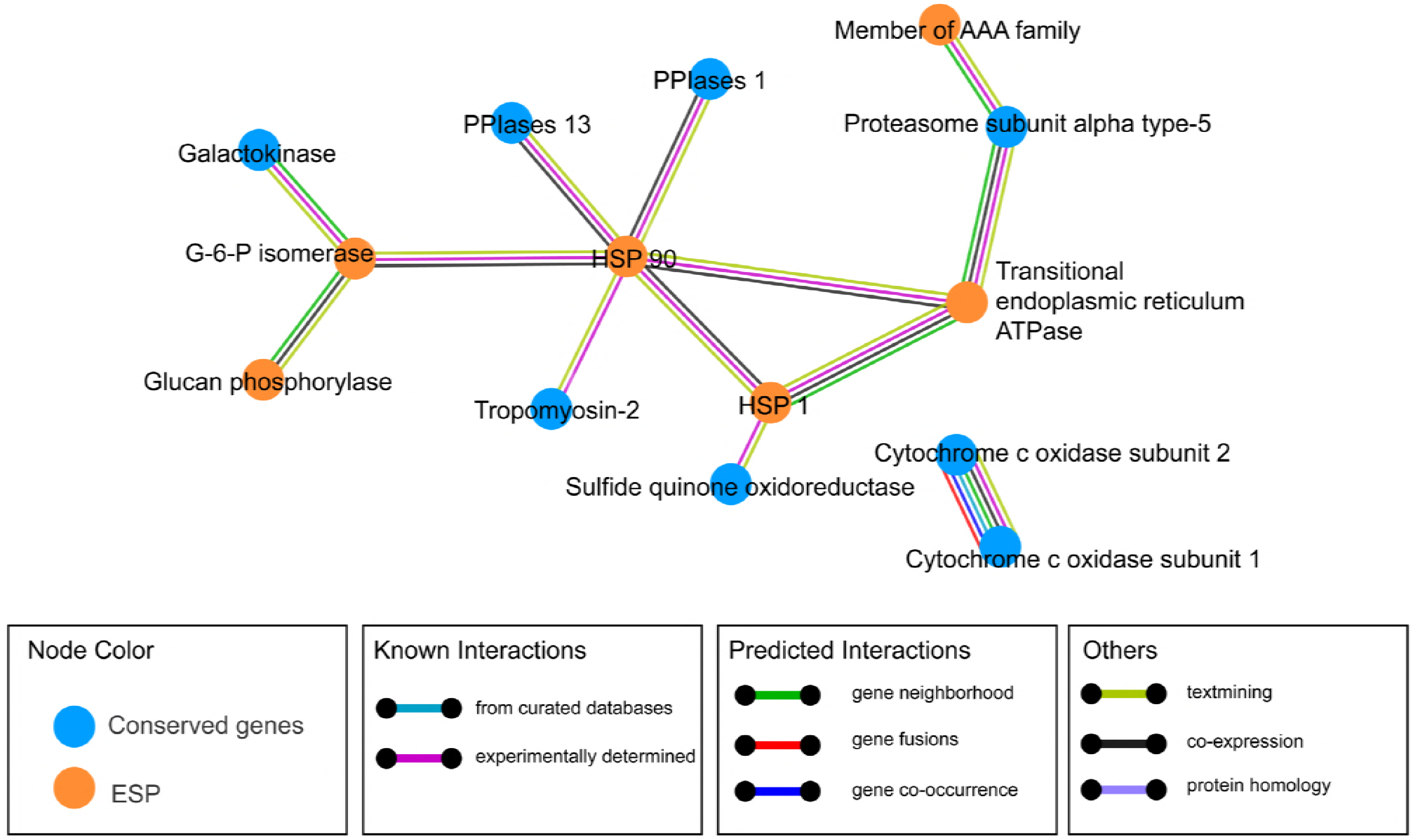
Protein-protein interactions of conserved genes and ESP genes.

## Discussion

The larval stage of the pork tapeworm *T. solium* is responsible for cysticercosis, which represents an important public health problem that occurs mainly in developing countries. *T. solium* cysticerci have developed diverse mechanisms to protect themselves from host immune attack [26], among which epigenetics may play an important role in gene regulation related to parasitism [24]. Recently, *Geyer* et al. found that essential DNA methylation machinery components, such as DNMT2 and MBD, are well conserved throughout the Platyhelminthes [4, 27]. Invertebrate DNMT2s are believed to retain strong DNA methyltransferase activity [28], which is different from vertebrate DNMT2s, which are considered tRNA methyltransferases [29]. Our computational searches indicated that both DNMT2 and DNMT3 are found in *T. solium*, which implies the potential existence of a more sophisticated DNA methylation machinery. In addition, *MBD2/3* homologs were also identified in the *T. solium* genome.

Based on these results, our present study focused on characterizing the DNA methylome and transcriptome of *T. solium* cysticerci, aiming for providing comprehensive omics profiles for this important parasitic stage of *T. solium*. We revealed a mosaic methylation pattern in *T. solium* that is typical of other invertebrates [3, 4, 21, 22, 30, 31]. Cytosine methylation was predominantly found in the CpA dinucleotide context, similar to other invertebrate species, including *Drosophila melanogaster* [32] and other *platyhelminths* such as *S. mansoni* [27], which might be mediated by MBD2/3 proteins [33, 34]. These patterns in the DNA methylome might be closely related to the activity of different DNMTs. As DNMT1 functions as a maintenance methylase by copying methylation after DNA replication with the help of Uhrf1 [35], a lack of DNMT1 might help to explain why much non-symmetrical methylation was observed in the *platyhelminth* genome. We also found that a periodicity for two pairs of mCpA and mCpT sites spaced with 13 bases between the pairs, corresponding to a single turn of the DNA helix, as previously observed. A structural study of the mammalian de novo methyltransferase DNMT3A and its partner protein DNMT3L found that two copies of each form a heterotetramer that contains two active sites separated by a length of 8-10 nucleotides in a DNA helix [36, 37]. Because we could not locate *DNMT3L* in the *T. solium* genome, the consistent 8-10 nucleotide spacing we observed in the *T. solium* genome might be due to *DNMT3A* alone or an unknown factor other than *DNMT3L*.

Gene methylation is believed to be an evolutionarily ancient means of transcriptional control. Among plants, vertebrates and some invertebrates such as *T. spiralis*, the notion that methylation in promoters primarily represses genes by impeding transcriptional initiation has been widely accepted [8, 18, 19], whereas intermediate levels of expression have been associated with genes experiencing the greatest extent of methylation in the gene body, indicating a bell-shaped relationship [38-40]. However, in invertebrates, such as the fungus *Neurospora crassa* [41] and the silkworm *Bombyx mori*, transcription initiation is unaffected. Thus, DNA methylation shows remarkable diversity in its extent and function across eukaryotic evolution. In our *T. solium* results, we also found that methylation levels of mCs were correlated with gene expression levels. Depending on different sequence contexts, methylation seemed to function differently in transcriptional regulation. Intriguingly, high CpG and CpC methylation levels of downstream regions, but not of promoter regions, were observed for silent genes (Figure 4). In contrast, upstream methylation seemed to mostly affect genes with low expression. Currently, knowledge on methylation patterns and their effects on gene regulation in non-vertebrates are still limited, though species-specific diversity has been observed [42]. Therefore, more data should be collected for the DNA methylomes of each specific species to characterize their patterns and functions.

In addition to characterizing the general distribution pattern of genome-wide DNA methylation, we also focused on methylation status of important *T. solium* genes. Based on previous studies, we looked into 27 extensively methylated genes that are important for *T. solium* development, survival and parasite-host interactions. We found 13 of these 27 genes mutually interacted based on annotations in the model organism *C. elegans*. Specifically, ESPs formed the core of the protein-protein-interaction network, while proteins encoded by conserved genes directly interacted with specific ESPs. Among the ESPs that were potentially regulated by DNA methylation, we found that two heat shock proteins (HSPs), hsp-90 and hsp-1, were highlighted and mutually interacted. The heat shock response is a general homeostatic mechanism that protects cells and organisms from the deleterious effects of environmental stress [43]. Together with COX-2, these proteins were previously reported to be important parasitism-related proteins [44]. Furthermore, we also revealed two genes encoding diagnostic antigens that might be regulated by DNA methylation, including diagnostic antigen gp50 and an 8 kDa diagnostic protein. GP50 is a glycosylated and GPI-anchored membrane protein. In recent years, one component of the lentil lectin purified glycoprotein (LLGP) antigens has been used for antibody-based diagnosis of cysticercosis [45]. The 8 kDa family members are metacestode excretory/secretory glycoproteins, which invoke strong antibody reactions in infected individuals [46]. Importantly, our data suggest that DNA methylation might play a key role in repressing their transcription, implying a potential for drug development in the future that can target epigenetic modification machinery to control this important neglected tropical disease.

## Acknowledgements

This study was supported by the National Natural Science Foundation of China (NSFC: 31460658, 31402185, 31440085, 31160504, 31030064, 31520103916) and the China Postdoctoral Science Foundation (2012M520674).

## Author contributions

M. Liu, F. Gao and S. Sun conceived and supervised the project. X. Liu and J. Wang performed NGS library construction. G. Ji and H. Lu conducted bioinformatic analysis. X. Bai, X. Wang, J. Pang, Y. Zhao, K. Yuan, X. Li collected and prepared samples. S. Sun prepared the manuscript. All authors have read and approved the manuscript for publication.

## Supplemental Materials

**Figures S1: Phylogenetic tree of DNMT proteins.**

**Figures S2: Phylogenetic tree of mbd proteins.**

**Figures S3: Patterns and chromosomal distribution of DNA methylation in *T. solium*.**

**Figures S4: DNA methylation patterns and chromosomal distribution.**

**Figures S5: Average density of methylation levels of cytosine distributed on genome.**

**Figures S6: Gene Ontology (GO) analysis for the genes with the lowest expression (2nd) and the most highly expressed (5th) genes.**

**Table S1: Results of reciprocal BlastP searches of T.s DNMTs and MBD.**

**Table S2: Results of ClustW on dnmt protein sequences.**

**Table S3: Data summary of MethylC-seq and RNA-seq.**

**Table S4: Summary of key parasitism genes that are methylated.**

## References

1. Schantz PM, Cruz M, Sarti E, Pawlowski Z: Potential eradicability of taeniasis and cysticercosis. Bull Pan Am Health Organ 1993, 27(4):397–403.

2. Sciutto E, Fragoso G, Fleury A, Laclette JP, Sotelo J, Aluja A, Vargas L, Larralde C: Taenia solium disease in humans and pigs: an ancient parasitosis disease rooted in developing countries and emerging as a major health problem of global dimensions. Microbes Infect 2000, 2(15):1875–1890.

3. Gao F, Liu X, Wu XP, Wang XL, Gong D, Lu H, Xia Y, Song Y, Wang J, Du J et al: Differential DNA methylation in discrete developmental stages of the parasitic nematode Trichinella spiralis. Genome Biol 2012, 13(10):R100.

4. Geyer KK, Rodriguez Lopez CM, Chalmers IW, Munshi SE, Truscott M, Heald J, Wilkinson MJ, Hoffmann KF: Cytosine methylation regulates oviposition in the pathogenic blood fluke Schistosoma mansoni. Nat Commun 2011, 2:424.

5. Aguilar-Diaz H, Bobes RJ, Carrero JC, Camacho-Carranza R, Cervantes C, Cevallos MA, Davila G, Rodriguez-Dorantes M, Escobedo G, Fernandez JL et al: The genome project of Taenia solium. Parasitol Int 2006, 55 Suppl:S127–130.

6. Lister R, Ecker JR: Finding the fifth base: genome-wide sequencing of cytosine methylation. Genome Res 2009, 19(6):959–966.

7. Feng S, Cokus SJ, Zhang X, Chen PY, Bostick M, Goll MG, Hetzel J, Jain J, Strauss SH, Halpern ME et al: Conservation and divergence of methylation patterning in plants and animals. Proc Natl Acad Sci U S A 2010, 107(19):8689–8694.

8. Zemach A, McDaniel IE, Silva P, Zilberman D: Genome-wide evolutionary analysis of eukaryotic DNA methylation. Science 2010, 328(5980):916–919.

9. Saitou N, Nei M: The neighbor-joining method: a new method for reconstructing phylogenetic trees. Mol Biol Evol 1987, 4(4):406–425.

10. Jones DT, Taylor WR, Thornton JM: The rapid generation of mutation data matrices from protein sequences. Comput Appl Biosci 1992, 8(3):275–282.

11. Kumar S, Stecher G, Tamura K: MEGA7: Molecular Evolutionary Genetics Analysis Version 7.0 for Bigger Datasets. Mol Biol Evol 2016, 33(7):1870–1874.

12. Trapnell C, Pachter L, Salzberg SL: TopHat: discovering splice junctions with RNA-Seq. Bioinformatics 2009, 25(9):1105–1111.

13. The Taenia solium Genome Project [http://www.taeniasolium.unam.mx/taenia/. Accessed 03 Sept 2016.]

14. Trapnell C, Williams BA, Pertea G, Mortazavi A, Kwan G, van Baren MJ, Salzberg SL, Wold BJ, Pachter L: Transcript assembly and quantification by RNA-Seq reveals unannotated transcripts and isoform switching during cell differentiation. Nat Biotechnol 2010, 28(5):511–515.

15. Tarailo-Graovac M, Chen N: Using RepeatMasker to identify repetitive elements in genomic sequences. Current protocols in bioinformatics 2009, 25:04.10.01–04.10.04.

16. Xi Y, Li W: BSMAP: whole genome bisulfite sequence MAPping program. BMC Bioinformatics 2009, 10:232.

17. STRING [http://string-db.org/. Accessed 03 Sept 2016.]

18. Zhang X: The epigenetic landscape of plants. Science 2008, 320(5875):489–492.

19. Weber M, Hellmann I, Stadler MB, Ramos L, Paabo S, Rebhan M, Schubeler D: Distribution, silencing potential and evolutionary impact of promoter DNA methylation in the human genome. Nature genetics 2007, 39(4):457–466.

20. Cokus SJ, Feng S, Zhang X, Chen Z, Merriman B, Haudenschild CD, Pradhan S, Nelson SF, Pellegrini M, Jacobsen SE: Shotgun bisulphite sequencing of the Arabidopsis genome reveals DNA methylation patterning. Nature 2008, 452(7184):215–219.

21. Xiang H, Zhu J, Chen Q, Dai F, Li X, Li M, Zhang H, Zhang G, Li D, Dong Y et al: Single base-resolution methylome of the silkworm reveals a sparse epigenomic map. Nat Biotechnol 2010, 28(5):516–520.

22. Lyko F, Foret S, Kucharski R, Wolf S, Falckenhayn C, Maleszka R: The honey bee epigenomes: differential methylation of brain DNA in queens and workers. PLoS Biol 2010, 8(11):e1000506.

23. Ye J, Fang L, Zheng H, Zhang Y, Chen J, Zhang Z, Wang J, Li S, Li R, Bolund L et al: WEGO: a web tool for plotting GO annotations. Nucleic Acids Res 2006, 34(Web Server issue):W293–297.

24. Robert McMaster W, Morrison CJ, Kobor MS: Epigenetics: A New Model for Intracellular Parasite-Host Cell Regulation. Trends in parasitology 2016, 32(7):515–521.

25. Victor B, Kanobana K, Gabriel S, Polman K, Deckers N, Dorny P, Deelder AM, Palmblad M: Proteomic analysis of Taenia solium metacestode excretion-secretion proteins. Proteomics 2012, 12(11):1860–1869.

26. Hewitson JP, Grainger JR, Maizels RM: Helminth immunoregulation: the role of parasite secreted proteins in modulating host immunity. Molecular and biochemical parasitology 2009, 167(1):1–11.

27. Geyer KK, Chalmers IW, Mackintosh N, Hirst JE, Geoghegan R, Badets M, Brophy PM, Brehm K, Hoffmann KF: Cytosine methylation is a conserved epigenetic feature found throughout the phylum Platyhelminthes. BMC Genomics 2013, 14:462.

28. Phalke S, Nickel O, Walluscheck D, Hortig F, Onorati MC, Reuter G: Retrotransposon silencing and telomere integrity in somatic cells of Drosophila depends on the cytosine-5 methyltransferase DNMT2. Nature genetics 2009, 41(6):696–702.

29. Goll MG, Kirpekar F, Maggert KA, Yoder JA, Hsieh CL, Zhang X, Golic KG, Jacobsen SE, Bestor TH: Methylation of tRNAAsp by the DNA methyltransferase homolog Dnmt2. Science 2006, 311(5759):395–398.

30. Bird AP, Taggart MH, Smith BA: Methylated and unmethylated DNA compartments in the sea urchin genome. Cell 1979, 17(4):889–901.

31. del Gaudio R, Di Giaimo R, Geraci G: Genome methylation of the marine annelid worm Chaetopterus variopedatus: methylation of a CpG in an expressed H1 histone gene. FEBS Lett 1997, 417(1):48–52.

32. Lyko F, Ramsahoye BH, Jaenisch R: DNA methylation in Drosophila melanogaster. Nature 2000, 408(6812):538–540.

33. Raddatz G, Guzzardo PM, Olova N, Fantappie MR, Rampp M, Schaefer M, Reik W, Hannon GJ, Lyko F: Dnmt2-dependent methylomes lack defined DNA methylation patterns. Proc Natl Acad Sci U S A 2013, 110(21):8627–8631.

34. Marhold J, Kramer K, Kremmer E, Lyko F: The Drosophila MBD2/3 protein mediates interactions between the MI-2 chromatin complex and CpT/A-methylated DNA. Development 2004, 131(24):6033–6039.

35. Law JA, Jacobsen SE: Establishing, maintaining and modifying DNA methylation patterns in plants and animals. Nat Rev Genet 2010, 11(3):204–220.

36. Huff JT, Zilberman D: Dnmt1-independent CG methylation contributes to nucleosome positioning in diverse eukaryotes. Cell 2014, 156(6):1286–1297.

37. Jia D, Jurkowska RZ, Zhang X, Jeltsch A, Cheng X: Structure of Dnmt3a bound to Dnmt3L suggests a model for de novo DNA methylation. Nature 2007, 449(7159):248–251.

38. Zilberman D, Gehring M, Tran RK, Ballinger T, Henikoff S: Genome-wide analysis of Arabidopsis thaliana DNA methylation uncovers an interdependence between methylation and transcription. Nature genetics 2007, 39(1):61–69.

39. Jjingo D, Conley AB, Yi SV, Lunyak VV, Jordan IK: On the presence and role of human gene-body DNA methylation. Oncotarget 2012, 3(4):462–474.

40. Nanty L, Carbajosa G, Heap GA, Ratnieks F, van Heel DA, Down TA, Rakyan VK: Comparative methylomics reveals gene-body H3K36me3 in Drosophila predicts DNA methylation and CpG landscapes in other invertebrates. Genome Res 2011, 21(11):1841–1850.

41. Rountree MR, Selker EU: DNA methylation inhibits elongation but not initiation of transcription in Neurospora crassa. Genes Dev 1997, 11(18):2383–2395.

42. Schubeler D: Function and information content of DNA methylation. Nature 2015, 517(7534):321–326.

43. Ferrer E, Gonzalez LM, Foster-Cuevas M, Cortez MM, Davila I, Rodriguez M, Sciutto E, Harrison LJ, Parkhouse RM, Garate T: Taenia solium: characterization of a small heat shock protein (Tsol-sHSP35.6) and its possible relevance to the diagnosis and pathogenesis of neurocysticercosis. Experimental parasitology 2005, 110(1):1–11.

44. Choi W, Chu J: The characteristics of the expression of heat shock proteins and COX-2 in the liver of hamsters infected with Clonorchis sinensis, and the change of endocrine hormones and cytokines. Folia parasitologica 2012, 59(4):255–263.

45. Hancock K, Pattabhi S, Greene RM, Yushak ML, Williams F, Khan A, Priest JW, Levine MZ, Tsang VC: Characterization and cloning of GP50, a Taenia solium antigen diagnostic for cysticercosis. Molecular and biochemical parasitology 2004, 133(1):115–124.

46. Ferrer E, Sanchez J, Milano A, Alvarez S, La Rosa R, Lares M, Gonzalez LM, Cortez MM, Davila I, Harrison LJ et al: Diagnostic epitope variability within Taenia solium 8 kDa antigen family: implications for cysticercosis immunodetection. Experimental parasitology 2012, 130(1):78–85.

